# Spatial heterogeneity in forest carbon storage affects priorities for reforestation

**DOI:** 10.1101/2021.07.06.450936

**Authors:** Rebecca Chaplin-Kramer, Justin Andrew Johnson, Richard P. Sharp, Julia Chatterton, Charlotte Weil, Alessandro Baccini, Sarah Sim

## Abstract

Reforestation is an important strategy for nature-based climate solutions and identifying carbon storage potential of different locations is critical to its success. Applying average carbon values from forest inventories ignores the spatial heterogeneity in forest carbon and the effects of forest edges on carbon storage degradation. Here we show how spatially-explicit, predictive carbon modeling, that leverages satellite, social and biogeophysical datasets, can be used to identify more efficient restoration opportunities for climate mitigation than area-based carbon stock averages. Accounting for regeneration of forest edges, in addition to reforestation, boosts estimates of potential carbon gains by more than 20%. The total potential carbon gain that could be achieved through reforestation at the level indicated by the Bonn Challenge (350Mha) is 51 Gt CO_2_-eq, but the “missing carbon” in our current forests accounts for 64.6 Gt CO_2_-eq globally; the greatest potential carbon gains are found in areas of high fragmentation.

## Main

Reforestation is touted as the largest natural pathway for climate mitigation,^1^ and forest-based mitigation comprises one-quarter of emission reductions planned by countries.^2^ Much recent effort has been devoted to identifying the lowest-cost restoration opportunities, offering the greatest carbon gain for the least or cheapest land area.^3,4^ A growing set of international initiatives promotes large scale reforestation to address climate change amidst other challenges like rural poverty and biodiversity loss. The Bonn Challenge has a restoration target of 350 Mha by 2030 and is already halfway toward securing national commitments,^5^ while under the UNFCCC^3^ countries have included more than 120 Mha of reforestation in their national climate pledges. The upcoming UN Decade on Restoration, along with the Trillion Trees Initiative that has garnered attention from public and private sectors alike,^6^ will continue to push forward financing and other incentives for reforestation.

While spatially-explicit global mapping of biomass carbon stocks using satellite technology is growing increasingly sophisticated,^7,8^ such data are only currently utilized for informing potential carbon loss from deforestation; assessment of the potential carbon gains through reforestation most commonly applies ecoregional or biome-level carbon stock averages.^1,3,9^ This misses the heterogeneity in carbon storage across forests due to biogeophysical variability or spatial configuration, which has been well documented in field^10,11^ and remotely-sensed observations.^12,13^ Forest edges contain up to 25% less carbon than their interiors^14^ and cause additional emissions of 0.34 Gt C per year or nearly a third of the emissions caused by deforestation.^15^ Without considering spatial heterogeneity in forest carbon, mitigation potential from reforestation could be vastly misrepresented and reforestation funding misallocated. Here we present a global, spatially-explicit, predictive model for carbon storage to evaluate carbon storage potential of forest restoration in different locations.

This global forest carbon model, based on a support vector machine (SVM) regression, explains 86% of the variability in above-ground biomass derived from satellite observations^7^ (Fig S1a, S2, Table S1) through climate, soil, topography, socio-economic factors, and land-use configuration. We compare the predicted carbon in above ground biomass resulting from this regression approach (Fig. S1b) to a more conventional approach to assessing carbon storage potential, based on mean values per land-use class in different regions or carbon zones (IPCC Tier 1 data^16^; Fig. S1c). We apply both approaches to a map of global reforestation potential, based on potential natural vegetation in forested biomes,^17^ excluding current agricultural and urban areas. Using the 350 Mha ambition of the Bonn Challenge^5^ as an area target, we optimize the selection of potential forest pixels to maximize carbon storage in ten 35 Mha increments. We also partition the gains in carbon into gains from restoration (i.e. converting non-forest back to forest) and gains from regeneration (increasing biomass in adjacent existing forest areas, through amelioration of edge effects). The difference in the results obtained from the regression versus the IPCC approach demonstrates how accounting for edge effects and spatial variability in forest biomass could influence landscape planning—in terms of both the location of optimal reforestation areas and the magnitude of potential carbon gain that may result.

### Prioritizing restoration areas

The two approaches produce dramatically different restoration priorities, only overlapping in 35% of their selected areas, mainly in Brazil and central Africa (Fig. 1). The regression approach prioritizes forests in the temperate as well as tropical regions, while the IPCC approach is focused almost exclusively in the tropics, and prioritizes a much larger area for restoration in Africa.

**Figure 1.**
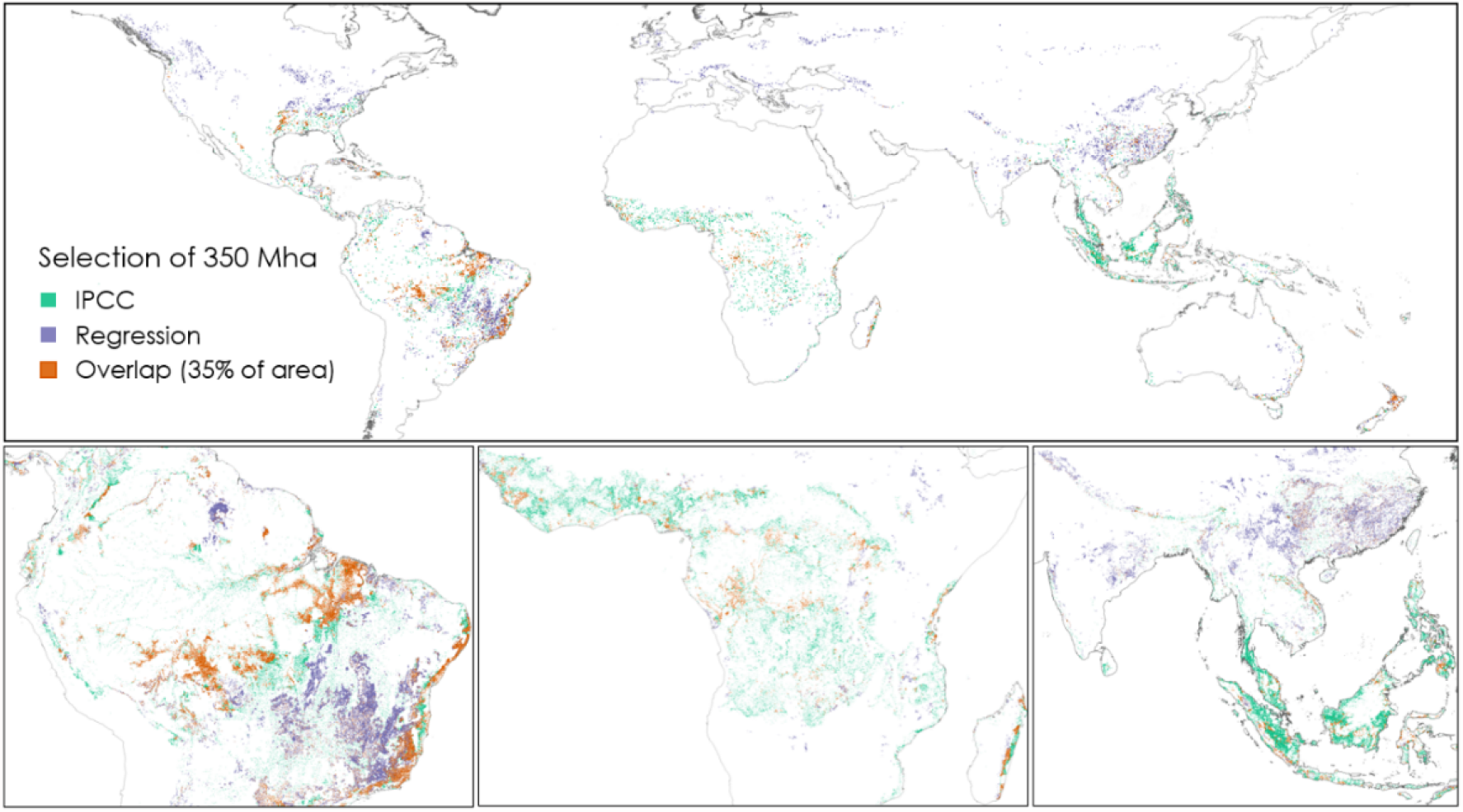
Restoration scenario for the Bonn Challenge (350 Mha), selected using the IPCC method (in green) or regression method (in purple) to optimize carbon storage potential. Overlaps between the two are shown in orange. Close-ups for the Amazon, the Congo, and Southeast Asia are shown in the inset.

The carbon gain in the reforestation area selected through the regression approach is up to 9.8 Gt CO_2_-eq higher than for the area selected through the IPCC approach, a difference accounting for nearly 20% of the total estimated carbon gain of 51.0 Gt CO_2_-eq for the 350 Mha reforestation target (Fig. 2a). The proportion of the total carbon gain resulting from regeneration (i.e., attributed to reduction of edge effects in existing forest, rather than addition of new forest) increases with the area of reforestation. Potential carbon gain in the forest edge reaches 24% of that in the new forest (Fig 2b). Comparisons with other restoration prioritization methods^9^ reveal that areas selected only by the regression (not overlapping with areas selected by IPCC) are at higher risk of further future degradation, and areas that are selected only by the IPCC method (not overlapping with areas selected by the regression) have slightly lower reforestation potential (in terms of difference between current and potential tree density; Fig. S4). This suggests that the regression method selects higher risk but also higher value areas for reforestation, which could benefit from policy intervention.

**Figure 2.**
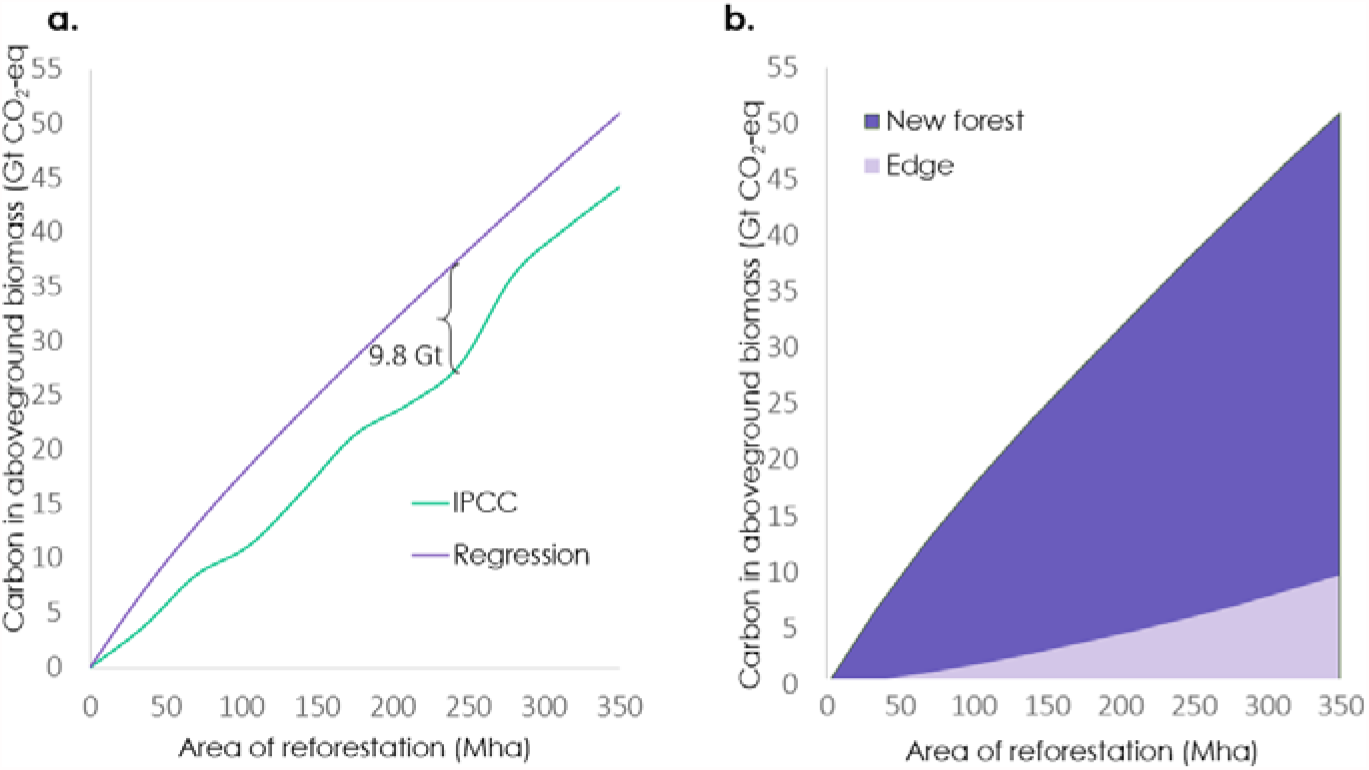
Carbon accumulation curve for IPCC method (green) and regression method (purple) for the total amount of aboveground biomass gain (a) and broken down into gain from new forest (dark purple) and forest edges (light purple) for the regression method (b). The total area of forest included in the restoration scenario (350 Mha) corresponds to the solution(s) shown in the map in Fig. 1. Carbon gain from reforestation selected using the regression method is up to 9.8 Gt CO_2_-eq greater than when selection occurs using the IPCC method. Gain within forest edge (due to reduction in edge effects from reforestation) is up to 24% of the gain from new forest.

The majority of lands selected by the regression for new forest are from crop mosaic, grassland and shrubland (Fig. 3a), with a fairly consistent proportion of degraded land within each class (with shrubland and grassland the highest at 27% and 23%, respectively). Globally, 22% of the land selected for restoration is classed as degraded (Fig. 3b), which is comparable to the proportion of degraded land globally (19%, according to the World Desertification Atlas^18^), suggesting that the optimization does not preferentially select degraded lands, though this varies considerably by country (Table S2).

**Figure 3.**
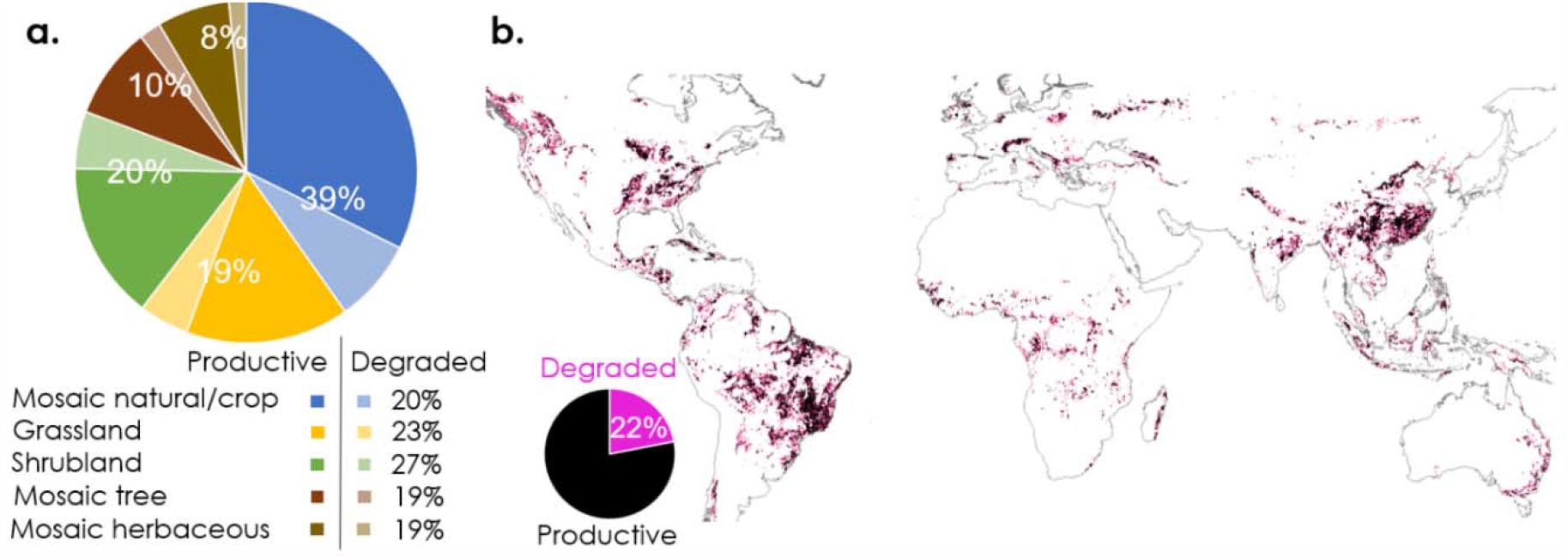
Existing land cover of areas selected for new forest in the regression-based optimization. a) Percentage of total lands selected by optimization, with productive lands in solid and degraded land in transparent shades; percentage of each class that is degraded shown in legend. b) Map of the 350 Mha selected for reforestation by the regression approach, with degraded lands in pink (comprising 22% of the total).

### Carbon gains possible in existing forest

The potential carbon gain that could be achieved through reforestation at the level indicated by the Bonn Challenge (51.0 Gt CO_2_-eq) is equivalent to a quarter to half of the total potential assessed by experts to be a plausible amount of carbon sequestration via nature-based climate solutions (avoided deforestation, reforestation, improved plantation management, and soil carbon sequestration) by the end of the century.^19^ However, this amount is considerably smaller than the potential gain if it were possible to eliminate all forest carbon edge effects (Fig 4). This “missing carbon” in our current forests accounts for 64.6 Gt CO_2_-eq globally, with the greatest potential gains found in areas of high fragmentation such as the Southeastern US, Central Africa, China, Europe, and parts of Southeast Asia and the Amazon. For the most part, these areas of greatest missing carbon occur in distinctly different regions from the areas selected for greatest gains through reforestation, with the exception of China.

**Figure 4.**
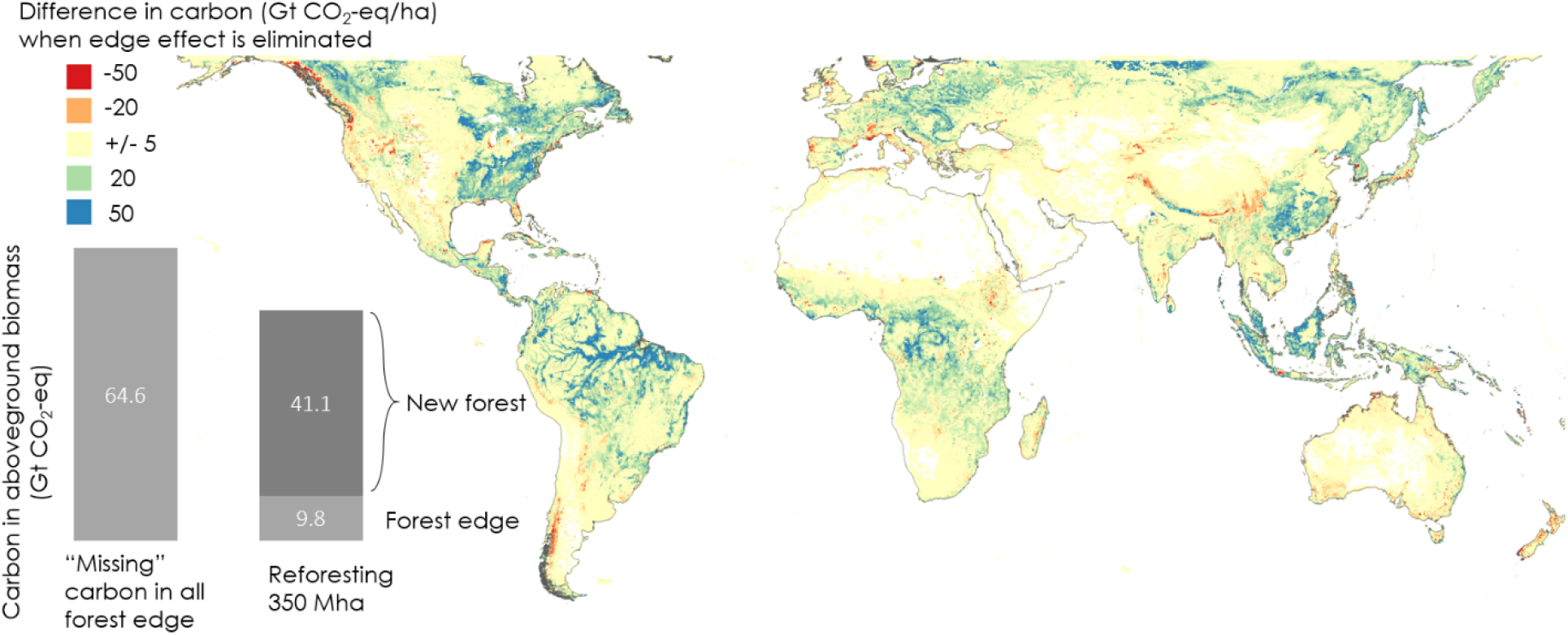
“Missing carbon” is the difference between carbon stored in aboveground biomass predicted by the regression model with the removal of edge effects (removing the influence of anthropogenic land-uses nearby) and that predicted under current conditions. Blue areas show where edge effects make the largest difference; orange and red areas show “reverse” edge effects, where predicted carbon is higher nearer to forest edges. Inset graph relates the total amount of “missing” carbon globally (64.6 Gt CO_2_-eq) to the potential carbon gain from reforesting at the level of the Bonn Challenge (51 Gt CO_2_-eq).

It is not realistic to expect to ameliorate the full extent of these edge effects everywhere, but the magnitude of this difference does indicate that addressing the possible causes of the “missing carbon” could provide a large source of carbon offsets, beyond conversion of non-forested lands back to forest. Modeling the “missing carbon” in fragmented and degraded forest areas could help identify where assisted regeneration through active management (e.g. seed dispersal or planting seedlings^20^) could have the greatest impact, or where it may be most important to address different human uses of forest causing degradation and carbon loss (e.g., selective logging, fuelwood harvest, and other activities that can turn a forest from a net sink to a net source of greenhouse gas emissions^21^). Further study with locally relevant landscape variables would be needed to identify the best strategies or interventions to replenish this “missing” carbon in different locations.

In addition to regions with “missing” carbon, there are also regions where edges appear to have a positive effect on carbon storage (denoted in red and oranges as negative “missing” carbon in Fig. 4), along the coast of British Columbia, in the Himalayan foothills, southern Andes, and parts of western Europe and eastern Africa. Such reversal in edge effects has been documented in field studies in temperate regions.^22^ In our analysis, these positive edge effect regions occur at different population densities, elevations and in different climates; the mechanisms behind higher carbon storage found in smaller forest fragments in these regions deserves further study.

Given the magnitude of these differences, the accuracy of the model and the underlying data are an important consideration. The results presented here are not biased toward the areas of highest error, either from the original Baccini dataset^7^ (Fig. S2) or empirical biomass data (Fig. S3). Biomass predicted by our regression is overestimated relative to Baccini throughout much of the Amazon and Africa, but this error does not consistently coincide with areas of greatest potential gains through reforestation (Fig 1) or regeneration (Fig 2). Biomass predicted by our regression is more often underestimated than overestimated relative to biomass found in forest plots (Table S1), which suggests the potential gains reported here are a conservative estimate. In general, the regression approach outperforms IPCC by more accurately representing patterns observed empirically (Fig. S3, Table S1), and thus we can have greater confidence in this spatially-explicit predictive approach.

### Potential policy implications

We have chosen the area outlined in the Bonn Challenge as a relevant policy target for which to demonstrate the utility of our new approach to carbon modelling, referencing international ambitions as currently articulated. However, we also explored a ‘total potential’ scenario of carbon storage gains from reforestation in any areas falling within forest biomes (with the exception of current agricultural and urban areas); our total potential reforestation extent (1074 Mha) would account for 119 Gt CO_2_-eq. This total potential area falls within the range of other estimates (345-1779 Mha estimated by Griscom et al.^1^; 900 Mha estimated by Bastin et al.^9^). Yet such comprehensive targets are not likely to be reached quickly, if at all, as even current commitments for the Bonn Challenge have not yet been met by the majority of countries that have reported their progress^5^. Thus, prioritization will be necessary, and the carbon gains achieved in the areas selected for reforestation in the early steps of the optimization using our regression approach are nearly double those achieved by the areas selected using the IPCC method (Fig. 2a).

Larger areas of restoration may be achievable through adequate carbon pricing (e.g., $20USD/tCO_2_eq could drive the restoration area to more than 800 Mha in total^3^), and economic analysis is an important next step for the exploration of forest carbon edge effects. Restoration costs, which are currently unavailable globally, would be a critical consideration when translating this global optimization to implementation at a landscape scale. However, moving beyond a focus on high-carbon-stock regions toward regions of high carbon storage potential through both reforestation and edge regeneration provides a first step to enabling more efficient allocations of restoration project resources.

Consideration of spatial variability including edge effects in forest carbon, and the potential for enhancing carbon gains through forest regeneration in addition to restoration, should be placed alongside other concerns that are emerging on reforestation, including multi-objective (for socio-economic as well as environmental objectives) planning and project permanence. Beyond reducing edge effects, there are many other reasons for prioritizing reforestation near forest edges, including greater likelihood of (lower-cost) natural, unassisted regeneration due to higher seed dispersal closer to forest remnants,^23^ and enhancing connectivity of fragments to enhance biodiversity.^24^ Evidence from programs already underway suggests that reforestation policies will fail to meet both biodiversity and carbon objectives if safeguards are not in place to prevent the degradation of one in pursuit of the other (as in the case of transitions to high-carbon plantations^25^). Restoration of degraded ecosystems, particularly in the Neotropics and Indomalaya, has been elevated as a key strategy to achieve these dual aims.^26^ Meanwhile, securing reforestation over the long term is becoming more challenging as climate change exacerbates threats that could lead to forest losses, such as fires, droughts, and pests or disease^27^; many of these same threats also magnify edge effects. While our current analysis does not include evaluation of restoration permanence, it is important to incorporate this into carbon offset programs, both for reforestation and edge regeneration, to avoid over-crediting.

The coming decade will determine whether and how an ambitious international agenda for climate action and sustainable development will be delivered. With many competing demands on our lands, we cannot afford to waste any opportunities to enhance carbon sequestration—properly accounting for spatial heterogeneity and forest edge effects can target investments for greatest benefit.

## Methods

### Regression vs. IPCC Tier 1 approach to predicting carbon storage

Our regression is based on the Baccini et al.^7^ (500 m) dataset, which includes a time series (2003-2014) to provide higher confidence in estimates due to an algorithm applied to remove outliers in individual years. Our regression is trained on the most recent year (2014), resampling the carbon data to the same resolution as the ESA LULC (10 arc sec, or ~300 m at the equator) used to define forest edge (see SI Appendix A2. Data). The global regression uses on-pixel predictors listed in SI Appendix A.1 and gaussian convolutions on LULC classes (non-forest, agriculture, and urban) to explore edge effects. It also includes interactions between edge and all other variables to account for the fact that edge effects vary across the different climates and soils represented in the different biomes. Although the aboveground biomass estimates made by Baccini et al. are themselves modelled, they are derived from very different sources than ours (a combination of reflectance, lidar, and field observations).

We compared this regression model to an IPCC Tier 1 model^16^ (Figure M1). The lookup-table method in the IPCC model is similar to a regression model, except that it only has a single value assigned to each LULC class (essentially, it is similar in concept to a regression based on nothing but dummy-variables for each LULC class). However, in reality, the IPCC method did not determine its coefficients from the regression, but rather from an exhaustive, literature-based analysis of observed carbon in specific carbon zones (the carbon zones in Ruesch and Gibbs are the cross-product of ecofloristic regions, frontier status and continent).

**Figure M1:**
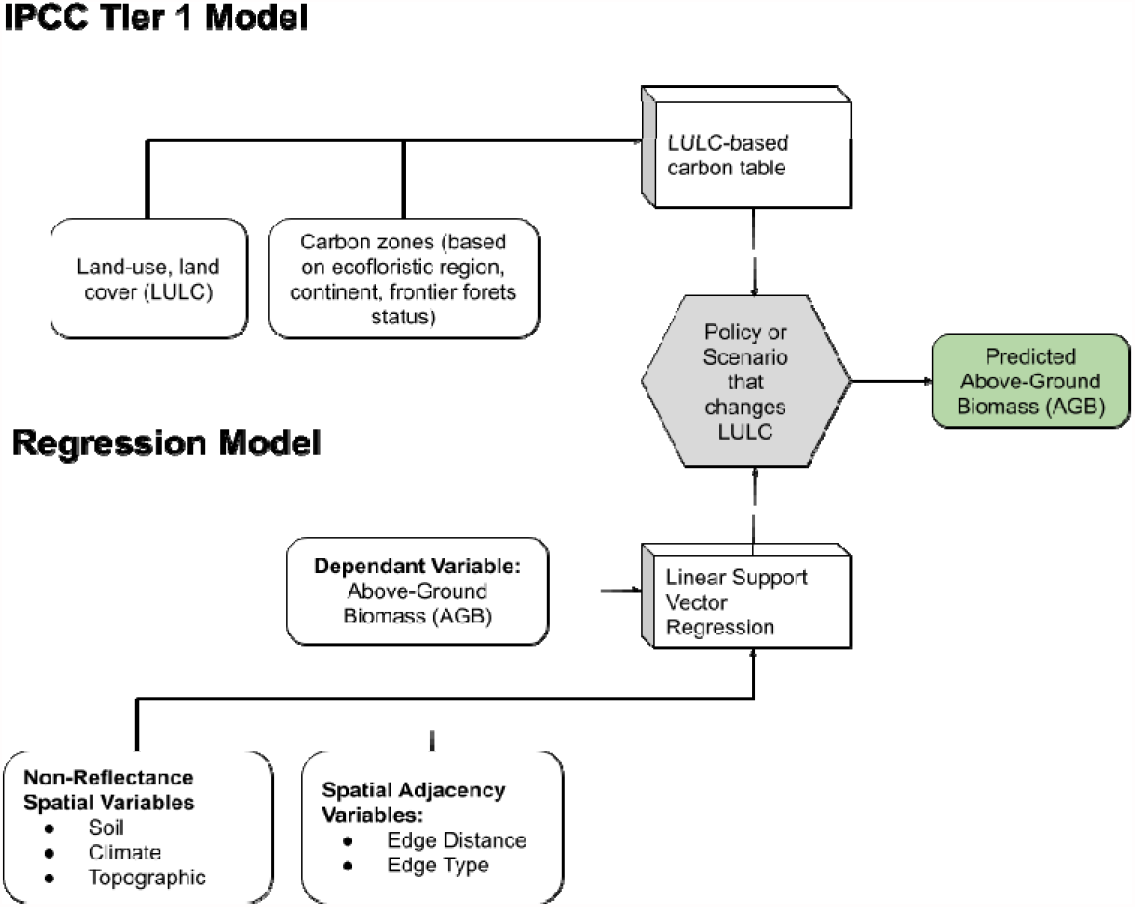
Modeling framework comparison between IPCC Tier 1 and Regression Model

We use Linear Support Vector Regression (LinearSVR) to address the problem of outliers which could dominate error in a simpler Ordinary Least Squares (OLS) regression. LinearSVR relaxes the optimization problem inherent in OLS to allow for a small amount of error in sample points away from the fit. In a 2D case, this error tolerance can be visualized as two lines (the support vectors) that bound the regression line. In multiple dimensions these are hyperplanes that bound the hyperplane regression. This method was chosen by comparing fits against an exhaustive search of other regression techniques that used cross-validation parameter searching techniques including OLS, Polynomial Regression, Ridge Regression, LARS-Lasso, Stochastic Gradient Descent, and Bayesian Regression. These searches were run across a subset of 90,000 sample points using an 80-20 random split for validation for each set.

### Linear Support Vector Machine Methodology

The LinearSVR technique fits a linear regression by solving the convex optimization problem that minimizes the L1 norm of the weights with the constraint that the error is less than a free “epsilon” error parameter. Formally, the following equation is minimized for the given samples:

where is the feature coefficient vector, is a constant value (set to 1 in our case), is the ith dependent variable, is the inner product of the feature coefficient vector with the ith dependent variable vector, *b* is the y-intercept, and is the free error parameter. The code to run this regression is available at: https://github.com/therealspring/carbon_edge_model/releases/tag/0.9.0 (model_builder.py).

### On-pixel effects

On-pixel effects are variables that explain the heterogeneity in carbon storage on the same pixel, including climate, soil, topographic, and socio-economic variables, of varying resolutions. See SI Appendix A.1.2 for a full list of these variables. We restrict our analysis here to forest only, but the same approach could be applied to any land use classes (though preliminary explorations not detailed here suggest the model fit is weaker outside of forest classes).

### Spatial adjacency (edge effects)

In order to explore edge effects on carbon storage, we also included spatial adjacency, variables for which nearby pixels explain heterogeneity in carbon storage. See SI Appendix A.1.3 for a list of these variables. These spatial adjacency predictor variables in our regression are assumed to have some relationship with the response variable (effect size), which depends on how great the distance is between the observed value of the independent (predictor) variable at location x and the resulting value of the dependent (response) variable at location y. When x = y, the coefficient is equivalent to that found when the on-pixel method is used. We allow the effect of these spatial adjacency variables (representing edge effects) to vary continuously over space, according to a flexible parametric equation based on the distance between any two grid-cells. Figure M2 illustrates this effect in one dimension and then in two dimensions.

**Figure M2.**
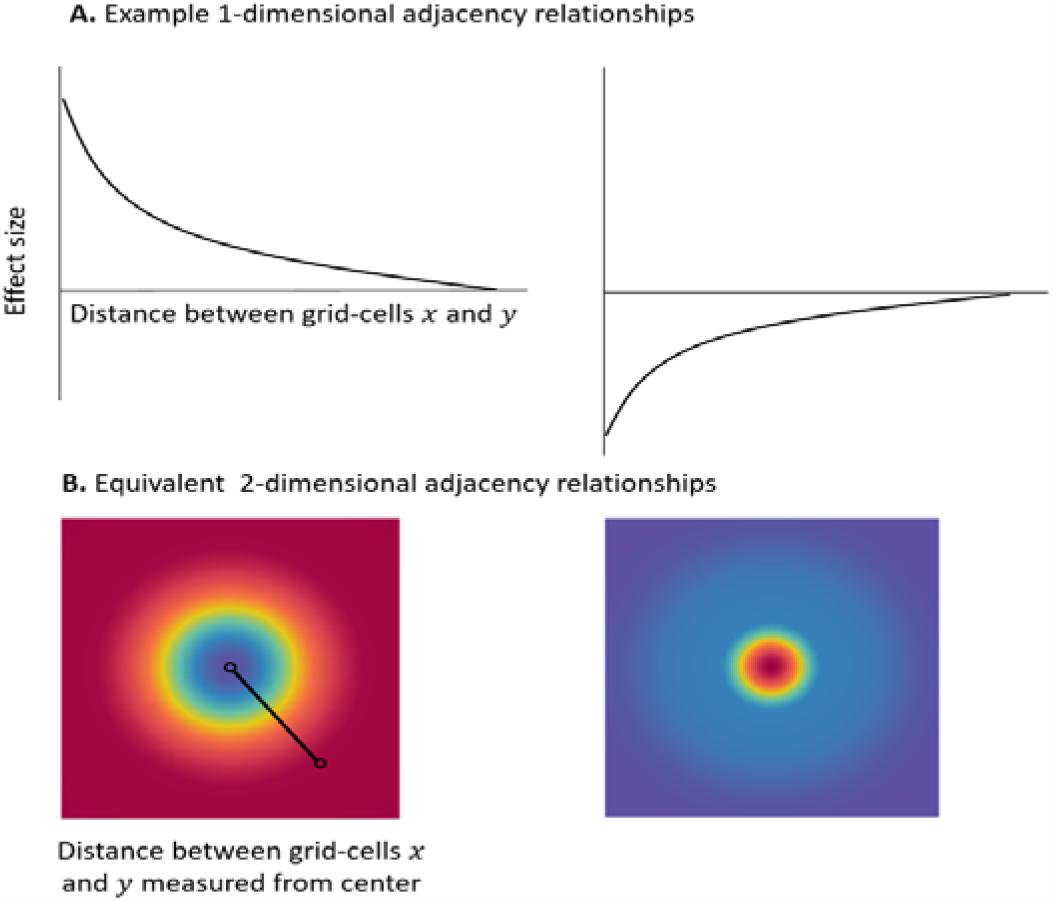
Adjacency functions and convolutions, demonstrating relationships for A) 1-dimensional adjacency (the distance between any two points) and B) 2-dimensional adjacency (with colors depicting the height or depth, warm/red colors denoting lower or more negative effects and the blue/cool colors denoting higher or more positive effects). The left side shows a strong positive effect of one pixel on another, declining as the distance between two points increases; the right side shows the inverse

After testing several different approaches to apply the general methods described above to our case studies, we concluded that using a range of Gaussian kernels in the regression framework was the best method. In practice, this means we calculate the Gaussian convolution of 0-1 LULC masks (binary values of 1 when the LULC is the class in question and 0 when it is not; e.g., 1 for forest, 0 for not-forest) for a variety of LULC classes using the following equation:

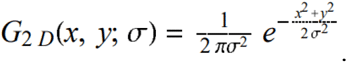

Given this, we choose a range of sigmas to use in this expression in order to generate different levels of smoothing. For example, a sigma of 4 means the effect of a variable on any given pixel will have half the strength of the effect on a pixel 4 pixels away (~1.2 km) and approximately zero effect on a pixel 10 pixels away (~3km in ESA data), according to a gaussian distribution. A lower value of sigma creates less smoothing while a higher value leads to more.

### Interactions and nonlinear effects

Edge effects differ across different regions, due to the influence of different variables that may exacerbate or ameliorate them. For example, greater anthropogenic pressures lead to stronger edge effects as people venture into the forest to harvest wood or graze livestock, while more moderate climates with regular temperatures lead to weaker edge effects because there is less biophysical stress at the forest edge. Therefore, in order to explore the impact of these different physical and socioeconomic variables on the resulting edge effects, we need to interact the convolution terms with every other predictor variable in the analysis. In order to determine the scale at which each variable has the greatest impact on edges, we interact each variable with all convolutions (from sigma 1-100; 300m -30 km). While there is the potential that multiple variables would interact together (for example, dimmer night lights may conceivably strengthen edge effects when population is high, indicating rural populations living more by wood power than electricity, but not when population is low, indicating a lack of human footprint altogether), the model complexity increases exponentially with the different combinations of variables, and directionality of three-way interactions is much more difficult to interpret. Therefore, we have left three-way interactions out of this analysis.

It is also possible that edge effects or other variables have nonlinear effects – exponential or saturating relationships with biomass. That is, they become disproportionately stronger at higher values (indicated by a positive coefficient for a second or third order term, the variable raised to the second or third power), or they reach a point after which continuing to increase has little effect (indicated by a negative coefficient for a second or third order term). We therefore include second and third order terms for all variables, but again, to reduce complexity from combinations, we don’t include second and third order terms of the interactions.

### Sub-sampling

As mentioned previously, we select a subset of 90,000 points across our input raster data to ensure that training with a LinearSVM method is computationally feasible. To generate any 2D raster point in that set, we first uniformly sample a 3D sphere then project that 3D point to a 2D coordinate using the same geographic projection the input raster data use for converting latitude/longitude coordinates to x/y coordinates on their 2D surfaces. This process of 3D to 2D sampling avoids biasing sample points against the geographic distortions that occur if the 2D surface were directly uniformly sampled. Once a point is selected, and if that point lies in a forest pixel, we use the data in the pixels in the raster stack under that point as a single data point for our regression training data. We continue this selection process until we have a desired number of points to train our model.

### Application of the global carbon regression model to restoration planning

Predictive carbon modeling is necessary for scenario assessment or landscape planning, to determine where and how much forest is needed to reach a carbon target, or how much additional carbon storage could be gained by reducing forest fragmentation. Here, we demonstrate the difference between our global regression model and a more typical approach based on IPCC Tier 1 data, informing how accounting for on-pixel carbon stock variability and edge effects would influence landscape planning. We identify priority areas for restoration under these two approaches through optimization, and assess the efficiency (how much carbon storage provided by how much area of restoration) for each. The optimization approximates the regression through a series of two-dimensional convolutions using gaussian kernels to select the pixels with the highest marginal value of returning to forest, by including not only the change in carbon on specific non-forest pixel, but also the distance-weighted potential for additional carbon from adjacent existing forest pixels whose edge effects would be reduced if that non-forest pixel is returned to forest (see Fig. M2). We also partition the gains in carbon into gains from restoration (i.e. converting non-forest back to forest) and gains from regeneration (increasing biomass in degraded forest areas, through amelioration of edge effects). The code to run these scenarios is available at https://github.com/therealspring/carbon_edge_model/releases/tag/0.9.0 (esa_restoration_optimizations.py).

We apply both the IPCC and the regression approaches to two scenarios: Current (based on ESA 2014) and a Restoration scenario. The Restoration scenario allows for the conversion of all habitat classified in a Potential Natural Vegetation (PNV) map as “forest” back to forest, with the exception of row crop agriculture (classes 10 and 20) and urban/residential (class 190). Classes allowed to be converted back to forest include pastures (appearing in ESA as grassland, class 130) and mosaics (both mosaic agriculture classes 30 and 40, and mosaic woody/herbaceous, classes 100 and 110), as well as any degraded “natural” land cover such as shrub or bare-ground, as long as they were classified as forest in the PNV map. The PNV map was developed by Smith et al. (unpublished; cross-walked from the Dinerstein et al. 2018 biome map to ESA LULC classes; SI Appendix A.3.1).

For the IPCC approach, we use the Tier 1 data (SI Appendix A.3.2) to create an IPCC-based Marginal Value Map, the difference between the carbon values for the ESA 2014 and the Restoration scenario map. For the regression approach, the global regression model (SI Appendix A.3.3) is applied to the forest pixels of Current (ESA 2014) and Restoration maps. The Baccini data are used for non-forest classes (a predictive model for the non-forest classes is not needed because the reforestation scenario only changes from non-forest to forest (a regression to predict carbon for the non-forest classes was attempted but the fit was very poor, likely because there is much more error in the EO data, which was developed based on tree allometry). The difference between the Restoration biomass and Current biomass maps creates the Regression-based Marginal Value Map.

The resulting biomass maps for both approaches are in Mg biomass/ha so to convert to CO_2_-equivalents per pixel, the following calculation is performed: [pixel area in Ha] * (15.9992*2+12.011)/12.011 * 0.47 (with the molecular weights as described for the IPCC approach, and the 0.47 biomass to carbon conversion ratio given by the IPCC).

The next step is to construct efficient restoration maps by selecting areas that contribute the most gain in carbon sequestration. For the IPCC approach this is straightforward; pixels are selected in rank order by the largest marginal value first until the area or carbon target is reached. For the regression approach, however, this is difficult to do from a per-pixel marginal value map because reforesting a single pixel not only sequesters carbon in that pixel, but indirectly through its neighbors with an edge effect. A ‘brute force’ approach could be to flip each pixel in the scenario on and off and evaluate the total carbon gain or loss by selecting that pixel. This approach is prohibitive for global scale high resolution maps and unnecessary. Our regression model captures the edge effect through a series of two-dimensional convolutions using gaussian kernels. These operations are “separable” meaning each new pixel of forest only adds to the carbon sequestration stocks of itself and its neighbors. We can “unroll” this effect by applying a 2d gaussian convolution to the marginal value map, yielding a raster whose values represent the extra carbon stocks gained by reforesting that pixel in the pixel itself as well as its neighbors.

Thus, we implement the following for a regression-based optimization:

1. Create a per-pixel marginal value map by subtracting the total carbon stocks of a future scenario from a base scenario.
2. Perform a normalized two-dimensional convolution with a gaussian kernel with a standard deviation at 3 kilometers on the marginal value map generated in (1). The result is a raster whose pixel values are a good approximation of how much carbon will be gained when that pixel is selected for reforestation.
3. Finally, create optimal reforestation maps by selecting potential new forest pixels from the raster generated in (2) in decreasing order. Selecting pixels in this order gives us a good approximation of the ‘best” pixels that contribute not only to the carbon stock in that pixel but the carbon edge effect gain from neighboring pixels decaying at the expected gaussian rate.

We illustrate this method in Figure M3. In Figure M3(a) we present an example marginal value map that has 4 possible pixels to select for reforestation marked by “NF” (new forest). Without accounting for edge effects, one might select the pixel with “20.0” units of potential carbon sequestration. Yet by processing that raster through a 2d gaussian convolution in (b) we get (c) which shows the most optimal pixel for reforestation is the upper left netting 34.7 units of carbon while the pixel selected in (a) would have only net 33.4.

**Figure M3.**
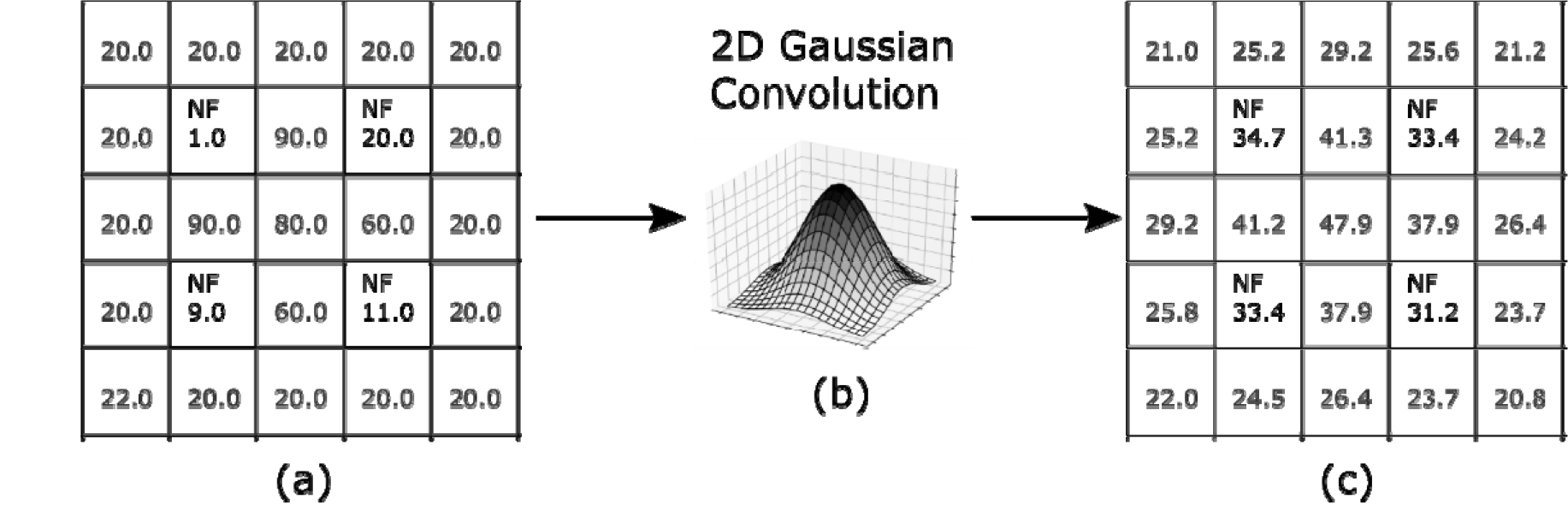
(a) Marginal value raster where NF are pixels of “new forest” that can be selected. Selecting them from max to min will give a suboptimal result because it does not account for forest edge effects. (b) Applying a 2D gaussian convolution (blur) to (a) yields (c). Note the upper left NF pixel will add an overall value of 34.7 units of carbon while selecting the “max” value in (a) will yield only 33.4.

## Supporting information

Appendix A - Supplemental info on data sources

## Acknowledgments

This work was funded by Unilever and a Stanford Data Science Initiative SEED grant. We are grateful to Jean-François Bastin, Lennart Olsson, Michael Cherlet, Katie Reytar, Peter Verburg, Jeff Smith and Holly Gibbs, Raphael Renevey, Alma Vexina Wilkinson, and Athanasios Nenes for their assistance in processing data.

## Author Contributions

RCK directed the work, performed post-hoc analyses of regression and optimization results, and drafted the manuscript; JAJ trained preliminary regression models, processed data, and drafted the methods; RPS gathered data, trained the final regression model, performed the optimization, and drafted the methods; CW gathered and processed data and performed exploratory analyses; AB provided and re-processed as necessary biomass and biomass uncertainty datasets and provided guidance in interpreting results; SS and JC evaluated many iterations of modeling outputs and SS set the scope of the problem for relevance to decision-making; all authors contributed to revision and improvement of the manuscript.

