## Supplemental figures and tables for "Spatial heterogeneity in forest carbon storage affects priorities for reforestation"

### Supplementary Figures and Tables

Figure S1. Aboveground biomass (ABG) predicted by (a) satellite data (Baccini), (b) regression, and (c) IPCC. Source: Baccini et al. 2017; Ruesch & Gibbs 2008.

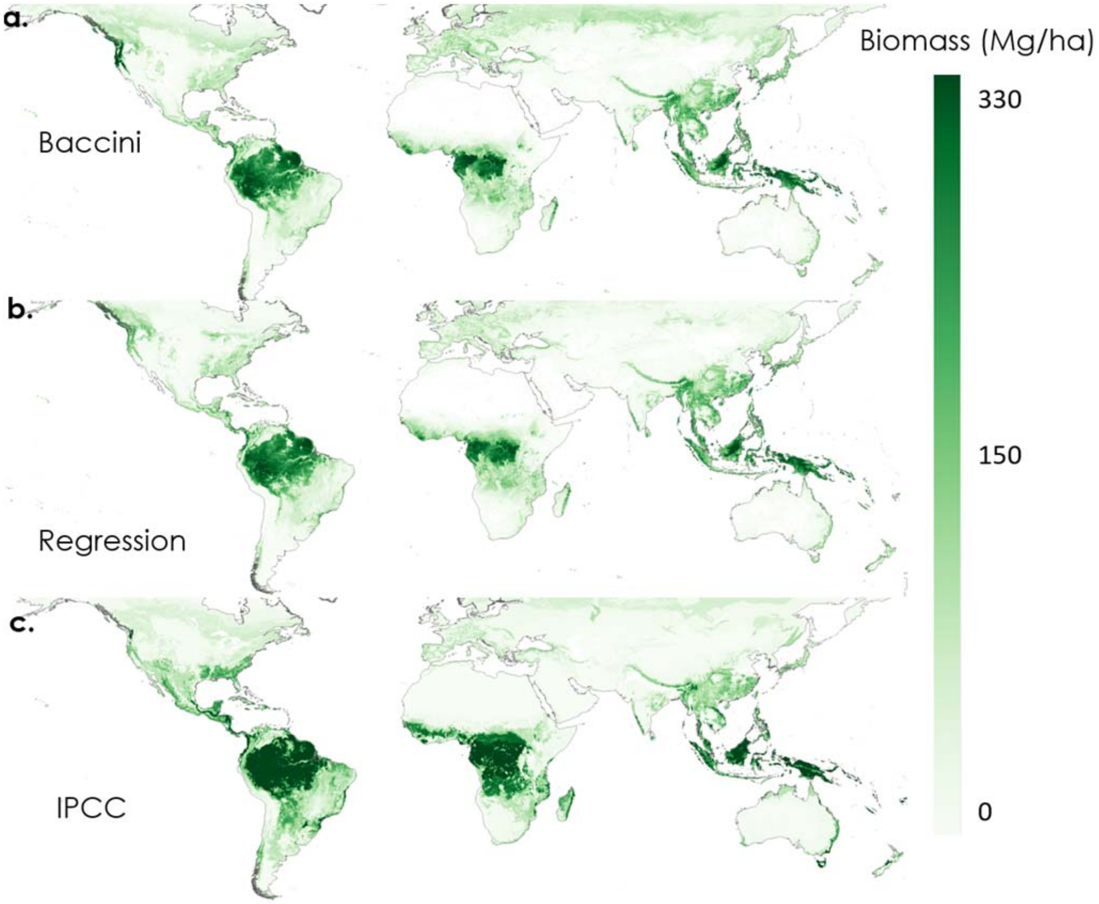

Figure S2. Regression model error relative to Baccini, showing areas where the model overestimates (in orange) and underestimates (in purple).

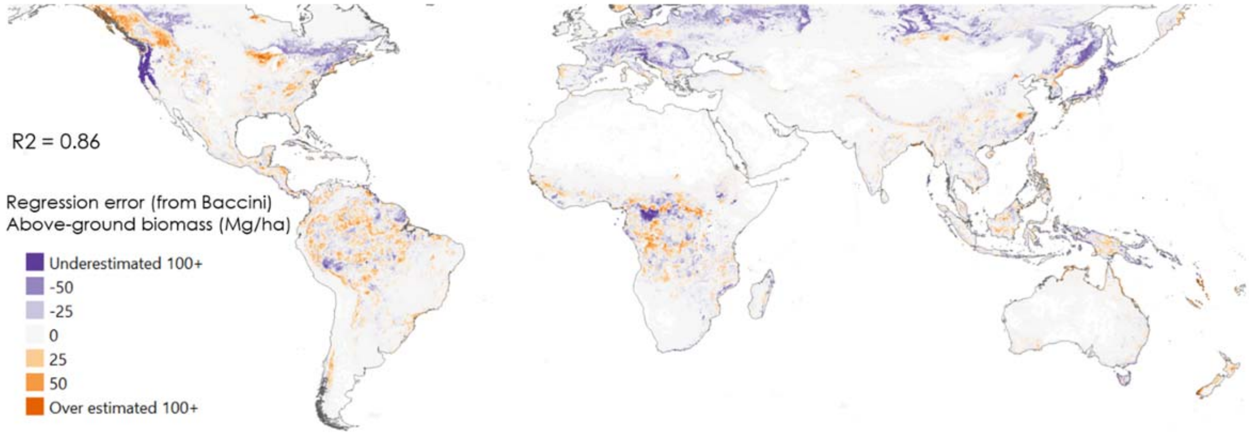


Figure S3. Accuracy of predictions from IPCC vs. regression approaches, relative to empirical biomass data from forest plots. Size of dots indicates magnitude of difference in the error of the two approaches, with red denoting plots where IPCC estimates were closer to forest plot data and blue denoting plots where the regression estimates were closer.

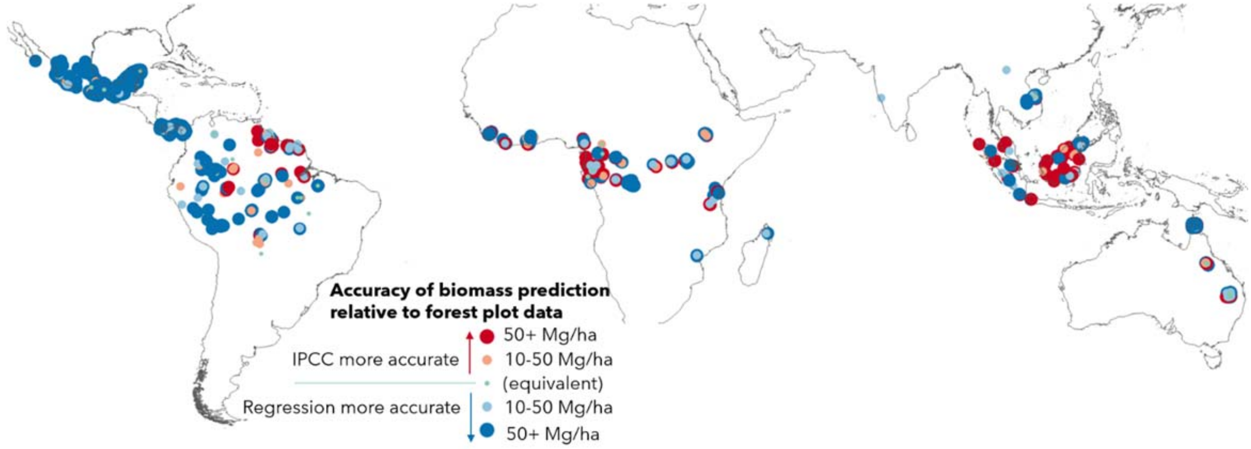

Fig. S4. Mean levels of future risk (projected change in tree cover; Fig. 3 in Bastin et al. 2019) and restoration potential (in terms of the difference between current and potential tree density; Fig 2B in Basin et al. 2019) for the areas selected by the IPCC method, regression method, and both (overlap; corresponding to Fig. 1). Future risk corresponds to negative values of projected change in tree cover, and the positive mean values for the areas selected by IPCC and the overlap between the two methods suggests that these areas are likely to regenerate on their own in the future. The slightly negative mean values for the areas selected by the regression method only suggests that these areas would require intervention for reforestation to occur. The lower mean values for restoration potential in areas selected by IPCC method only suggest that this method does not capture the highest carbon density possibilities. Source: Bastin et al. 2019.

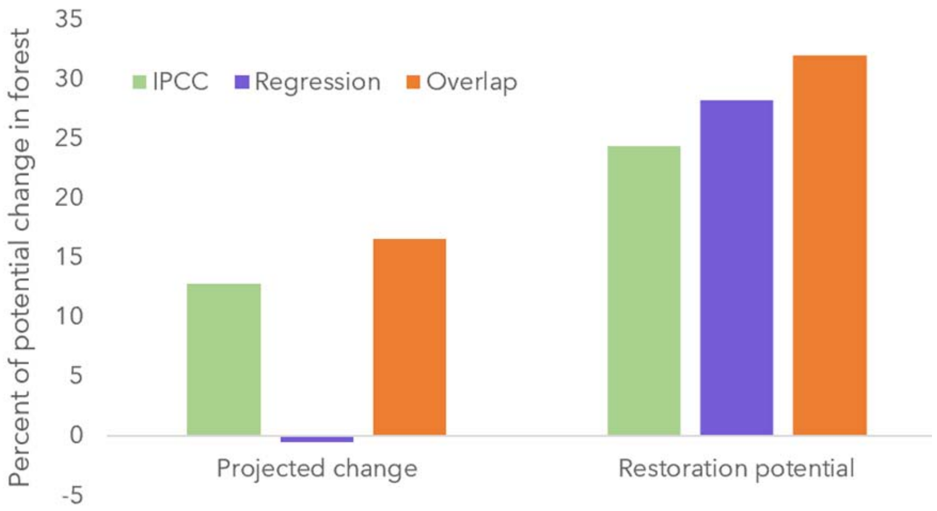

Table S1. Error in IPCC, regression and Baccini predictions of biomass, relative to empirical biomass data from forest plots.

| | **IPCC** | **Regression** | **Baccini** |
|---|---|---|---|
| RMSE (Mg/ha) | 155.3 | 114.5 | 110.8 |
| Avg % error | 91% | 12% | 15% |
| Avg overestimate | 134% | 77% | 72% |
| Avg underestimate | -30% | -34% | -33% |
| # overestimates | 74% | 41% | 45% |
| # underestimate | 26% | 59% | 55% |

Table S2. Area selected by the regression optimization in each country, and percent of degraded lands (selected by the optimization and in total), ranked by area selected, for countries with over 1 million hectares (Mha) selected.

| Country | Area selected (Mha) | Percent of country area selected | Total percent degraded | Percent degraded in 350Mha selection |
|---|---|---|---|---|
| Brazil | 118.98 | 14% | 31% | 25% |
| China | 43.23 | 5% | 6% | 8% |
| United States of America | 33.00 | 3% | 22% | 20% |
| Canada | 6.52 | 1% | 27% | 17% |
| India | 10.07 | 3% | 11% | 10% |
| Russia | 5.26 | 0% | 12% | 11% |
| Indonesia | 10.60 | 6% | 21% | 16% |
| Australia | 7.85 | 1% | 37% | 37% |
| New Zealand | 5.98 | 22% | 22% | 20% |
| Democratic Republic of the Congo | 5.19 | 2% | 15% | 7% |
| Bolivia | 4.84 | 4% | 26% | 17% |
| Mexico | 3.75 | 2% | 18% | 20% |
| Colombia | 4.04 | 4% | 15% | 13% |
| Vietnam | 3.28 | 10% | 34% | 32% |
| Madagascar | 3.04 | 5% | 17% | 34% |
| Myanmar | 2.87 | 4% | 24% | 22% |
| Venezuela | 2.97 | 3% | 19% | 14% |
| Paraguay | 2.75 | 7% | 48% | 49% |
| Argentina | 2.29 | 1% | 56% | 52% |
| United Kingdom | 1.41 | 6% | 18% | 16% |
| Italy | 1.68 | 6% | 9% | 11% |
| Cuba | 2.11 | 19% | 3% | 8% |
| Philippines | 2.19 | 7% | 10% | 14% |

| | | | | |
|---|---|---|---|---|
| Republic of the Congo | 2.23 | 6% | 12% | 9% |
| Nigeria | 1.99 | 2% | 31% | 31% |
| France | 1.45 | 2% | 9% | 12% |
| Poland | 1.07 | 3% | 1% | 4% |
| Spain | 1.30 | 3% | 10% | 17% |
| Honduras | 1.61 | 14% | 13% | 10% |
| United Republic of Tanzania | 1.58 | 2% | 32% | 33% |
| Papua New Guinea | 1.57 | 3% | 16% | 20% |
| Zambia | 1.54 | 2% | 6% | 7% |
| Peru | 1.51 | 1% | 27% | 13% |
| Chile | 1.18 | 2% | 17% | 33% |
| Haiti | 1.43 | 53% | 7% | 9% |
| Malaysia | 1.45 | 4% | 23% | 13% |
| Nicaragua | 1.41 | 11% | 11% | 10% |
| Guinea | 1.36 | 6% | 16% | 10% |
| Ethiopia | 1.35 | 1% | 36% | 30% |
| Pakistan | 1.07 | 1% | 7% | 8% |
