## Appendix A - Supplemental info on data sources for "Spatial heterogeneity in forest carbon storage affects priorities for reforestation"

### Appendix: Data sources for carbon analysis

#### A.1 Data sources for regression

##### A.1.1 Dependent variable (carbon datasets)

We use the Baccini et al. 2017 time series (2003-2014) of above-ground biomass to train our global regression. At 500 m resolution (which we resample to the 10 sec, ~300 m resolution of ESA LULC; see Supplementary Methods), it provides fine enough resolution to capture edge effects, but is more credible than any finer-resolution global products available. It has been validated and smoothed from year to year to avoid measurement error in individual years.[1]

##### A.1.2 Predictor variables, on-pixel

- WWF Ecoregions[2]
- Ecofloristic region[3]
- Topographic (as processed by [9])
    - Altitude
    - Slope
    - Aspect
    - Terrain Ruggedness Index
- Soil[4]
    - Grade of a sub-soil being acid e.g. having a pH < 5 and low BS
    - Available soil water capacity (volumetric fraction) with FC = pF 2.0
    - Available soil water capacity (volumetric fraction) with FC = pF 2.3
    - Available soil water capacity (volumetric fraction) with FC = pF 2.5
    - Depth to bedrock (R horizon) up to 200 cm
    - Probability of occurrence of R horizon
    - Absolute depth to bedrock
    - Bulk density (fine earth)
    - Cation Exchange Capacity of soil
    - Weight percentage of the clay particles (<0.0002 mm)
    - Volumetric percentage of coarse fragments (>2 mm)
    - Histosols probability cumulative
    - Soil organic carbon density
    - Soil organic carbon stock
    - Soil organic carbon content
    - pH index measured in water solution
    - pH index measured in KCl solution

---

[1] Baccini et al. 2017. Tropical forests are a net carbon source based on aboveground measurements of gain and loss. Science, 358(6360), 230-234

[2] Ricketts et al. 1999. Terrestrial ecoregions of North America: a conservation assessment.

[3] Ruesch, A., & Gibbs, H. K. (2008). New IPCC Tier-1 global biomass carbon map for the year 2000

[4] Hengl, T., Mendes de Jesus, J., Heuvelink, G. B.M., Ruiperez Gonzalez, M., Kilibarda, M. et al. (2017) SoilGrids250m: global gridded soil information based on Machine Learning. PLoS ONE 12(2): e0169748. doi:10.1371/journal.pone.0169748.

  - Sodic soil grade
  - Weight percentage of the silt particles (0.0002–0.05 mm)
  - Weight percentage of the sand particles (0.05–2 mm)
  - Keys to Soil Taxonomy suborders
  - World Reference Base legend
  - Texture class (USDA system)
  - Available soil water capacity (volumetric fraction) until wilting point
- Climatic[5]
  - Annual mean temperature
  - Annual mean precipitation
  - Subannual bioclimatic variables
    - Annual Mean Temperature
    - Mean Diurnal Range (Mean of monthly (max temp - min temp))
    - Isothermality (BIO2/BIO7) (* 100)
    - Temperature Seasonality (standard deviation *100)
    - Max Temperature of Warmest Month
    - Min Temperature of Coldest Month
    - Temperature Annual Range (BIO5-BIO6)
    - Mean Temperature of Wettest Quarter
    - Mean Temperature of Driest Quarter
    - Mean Temperature of Warmest Quarter
    - Mean Temperature of Coldest Quarter
    - Annual Precipitation
    - Precipitation of Wettest Month
    - Precipitation of Driest Month
    - Precipitation Seasonality (Coefficient of Variation)
    - Precipitation of Wettest Quarter
    - Precipitation of Driest Quarter
    - Precipitation of Warmest Quarter
    - Precipitation of Coldest Quarter
- Wind speed[6]
- ESACCI[7] forest categorization -
  - 'Mosaic natural vegetation (tree, shrub, herbaceous cover) (>50%) / cropland(<50%)'
  - 'Tree cover, broadleaved, evergreen, closed to open (>15%)'
  - 'Tree cover, broadleaved, deciduous, closed to open (>15%)'
  - 'Tree cover, broadleaved, deciduous, closed (>40%)'
  - 'Tree cover, broadleaved, deciduous, open (15-40%)'
  - 'Tree cover, needleleaved, evergreen, closed to open (>15%)'
  - 'Tree cover, needleleaved, evergreen, closed to open (>15%)'
  - 'Tree cover, needleleaved, evergreen, open (15-40%)'

---

[5] Fick, S.E. and R.J. Hijmans, 2017. Worldclim 2: New 1-km spatial resolution climate surfaces for global land areas. International Journal of Climatology

[6] Global Wind Atlas. https://globalwindatlas.info/api/gis/global/wind-speed/50

[7] http://maps.elie.ucl.ac.be/CCI/viewer/

    - 'Tree cover, needleleaved, deciduous, closed to open (>15%)'
    - 'Tree cover, needleleaved, deciduous, closed (>40%)'
    - 'Tree cover, needleleaved, deciduous, open (15-40%)'
    - 'Tree cover, mixed leaf type (broadleaved and needle-leaved)'
    - 'Mosaic tree and shrub (>50%) / herbaceous cover (<50%)'
    - 'Sparse vegetation (tree, shrub, herbaceous cover) (<15%)'
    - 'Sparse tree (<15%)'
    - 'Tree cover, flooded, fresh or brackish water'
    - 'Tree cover, flooded, saline water'
- - ESACCI Non-forest classes (0, 11, 12, 20, 30, 40, 100, 110, 120, 121, 122, 130, 140, 150, 151, 152, 153, 180, 190, 200, 201, 202, 210, 220)
    - 'Cropland, rainfed'
    - 'Cropland, rainfed, herbaceous cover'
    - 'Cropland, rainfed, tree or shrub cover'
    - 'Cropland, irrigated or post-flooding'
    - 'Mosaic cropland (>50%) / natural vegetation (tree, shrub, herbaceous cover)(<50%)'
    - 'Shrub or herbaceous cover, flooded, fresh/saline/brackish water'
    - 'Urban areas'
    - 'Bare areas'
    - 'Consolidated bare areas'
    - 'Unconsolidated bare areas'
    - 'Sparse shrub (<15%)'
    - 'Sparse herbaceous cover (<15%)'
    - 'Mosaic herbaceous cover (>50%) / tree and shrub (<50%)'
    - 'Shrubland'
    - 'Evergreen shrubland'
    - 'Deciduous shrubland '
    - 'Grassland'
    - 'Lichens and mosses'

##### A.1.3 Predictor variables, spatial adjacency

- Convolution (gaussian kernel) for ESA non-forest (see above)
- Convolution (gaussian kernel) for ESA agriculture (classes 10, 20, 30, 40)
- Convolution (gaussian kernel) for ESA urban (class 190)
- Convolution (gaussian kernel) for Settlements[8] (not available globally; for Baccini 30 m analysis only)
- Convolution (gaussian kernel) for tree cover[9] (at 30 m; for Baccini 30 m analysis only)
- Population density[10] (at 1km and 10 km)
- Market accessibility[11] (at 10 km)

---

[8] CIESEN/Facebook High Resolution Settlement Layer. https://www.ciesin.columbia.edu/data/hrsl/

[9] Global Forest Watch, Hansen tree cover product.

[10] Gridded Population of the World. Version 4. https://sedac.ciesin.columbia.edu/data/collection/gpw-v4

[11] Weiss et al. 2018. A global map of travel time to cities to assess inequalities in accessibility in 2015. Nature 553:333-336.

- Livestock density[12] (at 10 km)
- Night lights[13] (at 1km and 10km)

#### A.2 Data generated by the regression and used in subsequent analysis

- Model coefficients of selected model: https://storage.googleapis.com/ecoshard-root/global_carbon_regression/lasso_interacted_not_forest_gs1to100_nonlinear_alpha0-0001_params_namefix.csv
- Full set of results for all tested models for the global regression: https://storage.googleapis.com/ecoshard-root/global_carbon_regression/model_results_global_regression.zip
- Training and testing datasets for Baccini aboveground biomass datasets resampled to 10 s): https://storage.googleapis.com/ecoshard-root/global_carbon_regression/training_testing_subsample_baccini_10s.zip
- Forest plot points: https://storage.googleapis.com/ecoshard-root/global_carbon_regression/merged_forest_plot_data.gpkg
- Validation analysis: https://storage.googleapis.com/ecoshard-root/global_carbon_regression/regression_validation_on_forest_plots.xlsx

#### A.3 Data used in the scenario application of regression model

##### A.3.1 Data used for the restoration scenarios

- ESA LULC map[14]: https://storage.googleapis.com/ecoshard-root/global_carbon_regression/ESACCI-LC-L4-LCCS-Map-300m-P1Y-2014-v2.0.7_smooth_compressed.tif
- Restoration scenario (based on PNV, below, but excluding agriculture and urban): https://storage.googleapis.com/nci-ecoshards/scenarios050420/restoration_limited_md5_372bdfd9ffaf810b5f68ddeb4704f48f.tif
- Potential Natural Vegetation (PNV) full reforestation scenario: https://storage.googleapis.com/ecoshard-root/global_carbon_regression/PNV_jsmith_060420_md5_8dd464e0e23fefaaabe52e44aa296330.tif
- The optimization can be run through the esa_restoration_optimization.py script contained in the version of the repository archived at: https://zenodo.org/record/5056934#.YN95w-hKg2x

[12] Gridded Livestock of the World. Version 3. https://dataverse.harvard.edu/dataverse/glw_3
[13] DMSP-OLS Nighttime Light Series 1992-2013 https://ngdc.noaa.gov/eog/dmsp/downloadV4composites.html
[14] European Space Agency CCI Land Cover Project. http://maps.elie.ucl.ac.be/CCI/viewer/download.php

##### A.3.2 Data used for IPCC approach

- IPCC Tier 1 data[15] (carbon zones are rows and ESA LULC codes are columns): https://storage.googleapis.com/ecoshard-root/global_carbon_regression/IPCC_carbon_table_md5_a91f7ade46871575861005764d85cfa7.csv
- Carbon zones map: https://storage.googleapis.com/ecoshard-root/global_carbon_regression/carbon_zones_md5_aa16830f64d1ef66ebdf2552fb8a9c0d.gpkg

##### A.3.3 Data used for the regression approach

- All data needed to run the regression approach are contained within the model produced by the carbon_edge_model.py script found in the version of the repository archived at: https://zenodo.org/record/5056934#.YN95w-hKg2x
- The complete set of coefficients are available at: https://github.com/therealspring/carbon_edge_model/blob/master/images/coef_640000.csv

---

[15] Based on Ruesch & Gibbs 2008. Available online from the Carbon Dioxide Information Analysis Center [http://cdiac.ess-dive.lbl.gov], Oak Ridge National Laboratory, Oak Ridge, Tennessee.
